# Inter-layer and inter-subject variability of circadian gene expression in human skin

**DOI:** 10.1101/2022.06.03.494693

**Authors:** Marta del Olmo, Florian Spörl, Sandra Korge, Karsten Jürchott, Matthias Felten, Astrid Grudziecki, Jan de Zeeuw, Claudia Nowozin, Hendrik Reuter, Thomas Blatt, Hanspeter Herzel, Dieter Kunz, Achim Kramer, Bharath Ananthasubramaniam

## Abstract

The skin is the largest human organ with a circadian clock that regulates its function. Although circadian rhythms in specific functions are known, rhythms in the proximal clock output, gene expression, in human skin have not been thoroughly explored. This work reports circadian gene expression in two skin layers, epidermis and dermis, in a cohort of young, healthy adults, who maintained natural, regular sleep schedules. 10% of the expressed genes showed rhythms at the population level, of which only a third differed between the two layers. Broadly, expression magnitudes of circadian genes were consistent across subjects in each layer. Amplitude and phases of circadian gene expression, however, varied more across subjects than layers, with amplitude being more variable than phases. Expression amplitudes in the epidermis were larger and more subject-variable, while they were smaller and more consistent in the dermis. Core clock gene expression was similar across layers at the population-level, but were heterogeneous in the their variability across subjects. We used this data to identify small sets of biomarkers for internal clock phase in each layer, which consisted of layer-specific non-core clock genes. This work provides a valuable resource to advance our understanding of human skin to realize the potential of circadian medicine as well as a novel methodology to quantify sources of variability in human circadian rhythms.

## I. Introduction

The skin is the largest organ of the body and its main functions are protection against bacteria, radiation or temperature from the exterior, as well as against water loss from the interior [1]. It is morphologically complex and consists of many cell types that are organized into three main layers: epidermis, dermis and hypodermis [2]. The skin evolved a circadian clock [3] in response to the direct exposure to the rhythmic external environment to anticipate changes and to adjust its physiology accordingly.

The mammalian circadian clock is a hierarchical network with the central clock in the suprachiasmatic nucleus (SCN) and peripheral clocks in many tissues including the skin. The cell-autonomous molecular “core” clock [4, 5] consists of a number of interlocked transcriptional-translational negative feedback loops. Core clock genes *CLOCK* and *BMAL1* induce the expression of their own inhibitors, *PER* and *CRY* genes. Once translated, PER and CRY proteins form large complexes that travel back to the nucleus to repress CLOCK and BMAL1, thus repressing their own transcription and thereby creating self-sustained 24 h rhythms in gene and protein expression. In mammals, core clock components that are also transcription factors act at cis-regulatory sequences to drive rhythmic expression of a large number of output genes (about 10% of all genes) in a cell-autonomous and tissue-specific manner [6, 7].

The circadian gene expression profile of the skin remains nevertheless incompletely characterized. The presence of a skin circadian clock in humans was first inferred from circadian rhythms in biophysical skin parameters, such as sebum secretion [8], water loss [9] or response to allergens [10]. At the turn of the century, rhythmic expression of selected core clock genes in the skin was described in humans [11] and mice [12]. Over time, circadian expression of core clock genes was recorded in several skin cell types, including epidermal and hair follicle keratinocytes, dermal fibroblasts and melanocytes [13–17]. Spörl et al. [18] performed the first high-throughput analysis of circadian gene expression in the skin. That study identified ~300 circadian genes from measurements at three time points in one layer (epidermis). More recently, Wu et al. [19, 20] identified ~100-150 circadian genes each in the epidermis and dermis from samples collected every 6h over one day. However, both these microarray studies provide only limited insight into circadian gene expression in human skin, since they lacked sufficient number of samples over one circadian cycle. To more thoroughly describe the impact of the human skin clock, we identified circadian genes in the two prominent skin layers, epidermis and dermis. To assess whether the complex and heterogeneous skin also results in a cell type-/layer-specific clock, we compared the circadian gene expression across layers.

As one of the few accessible tissue clocks, skin samples could be used for circadian phenotyping of humans. Evidence is gradually accumulating that therapeutic efficacy and the degree of side effects are dependent on the time of administration [21–23]. Such observations are likely to grow, since 50% of all drugs target the product of a circadian gene [6, 7]. One key challenge to implementing time-of-day-aware ‘circadian medicine’ strategies is the fact that internal clocks of humans are heterogeneous. Since rhythms in human physiology are determined by *internal* clock time and not on time according to the external environment, circadian studies in humans ought to record and present results relative to the internal phase of entrainment (termed chronotype) of subjects. This internal clock time in turn depends on genetic factors [16,24], age [25], sex [25], level of light exposure [26, 27], the season [26, 28] and on the local time-zone [25]. Thus, circadian treatments need to be personalized to an subjects’s clock. To evaluate the utility of skin samples for circadian phenotyping, we used our comprehensive gene expression profiles to identify biomarkers for circadian phase in each layer.

We measured gene expression using microarrays in a small cohort of young, healthy adults of both sexes, who maintained their natural yet regular sleep schedules. We first quantified the circadian expression expected in the general population in the epidermis and dermis. In contrast to previous studies on skin, we present our results with respect to internal time of subjects; our study design included chronotyping of subjects. We then analyzed the layer-specificity of *population* circadian rhythms. The population circadian rhythms are the most representative rhythms of an individual in the population. However, the inherent heterogeneity will result in individual rhythms diverging from the population rhythm. Therefore, we next quantified how much individual circadian rhythms deviate from the population rhythms in both layers. Finally, we identified a small set of biomarkers with circadian gene expression to predict internal clock phase in each layer.

## II. Results

### Population circadian gene expression is layer-specific in healthy human skin

To explore molecular circadian rhythms in human dermis and epidermis, 11 healthy subjects (male and female) were biopsied in the upper back every 4 h across a 24 h duration (Figure 1A, Materials and Methods). In the two weeks leading up to the biopsies, subjects maintained their desired natural sleep-wake schedule. Samples were separated into dermis and epidermis and subsequently quantified using whole-genome microarrays. We adjusted sample collection times using their chronotypes to indicate *internal* time of subjects (Figure 1B). Chronotypes were estimated from sleep schedules (available in Supplementary Table S1) as the mid-sleep time on free days after correcting for sleep debt (MSF_sc_) [29].

**Figure 1:**
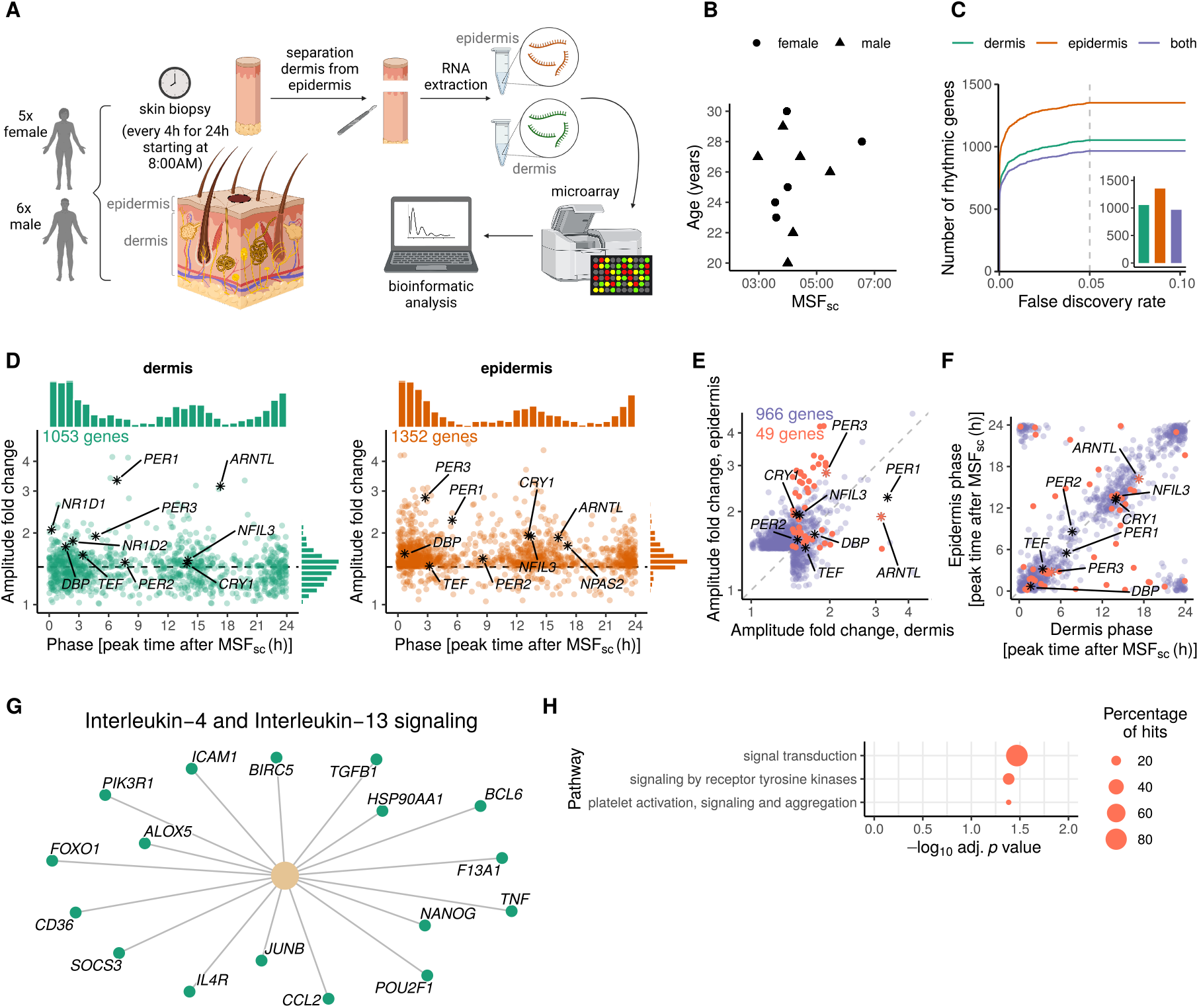
Functional and layer-specific clocks in human dermis and epidermis. **A.** Experimental setup: 11 healthy subjects were biopsied in the back every 4 h for 24 h starting at 8 AM. Dermis and epidermis were separated and gene expression was analyzed using whole-genome microarrays. **B.** Composition of the study cohort by sex, age and mid-sleep time on free days after correcting for sleep debt (MSF_sc_). **C.** Number of circadian genes with respect to *internal* time in dermis, epidermis or in both layers were determined using differential rhythmicity analysis (minimum requirement of peak-to-trough fold change amplitude >1.5 in at least one layer, see Materials and Methods for details). The number of genes for FDR =0.05 is shown in the inset. **D.** Phase (as peak time after MSF_sc_) and amplitude distributions of the circadian genes in human dermis (in green, left panel) and epidermis (orange, right panel) (FDR <0.05, peak-to-trough fold change amplitude >1.5). Each gene is represented by a dot; clock genes are highlighted in black. **E.** Amplitude correlation of the 966 circadian genes in both layers, from which 49 show significant different rhythms (highlighted in coral). **F.** Phase correlation of circadian genes in dermis *and* epidermis, with differentially rhythmic genes highlighted in coral. Clock genes are shown in black asterisks (or coral if differentially rhythmic). **G.** Pathway enrichment analysis of circadian genes in dermis was performed with REACTOME pathway database. Rhythmic genes belonging to the pathway are shown. **H.** Reactome pathway enrichment analysis of differentially-rhythmic genes in dermis and epidermis tested against the background of all 1439 rhythmic genes in dermis or epidermis. Only pathways with a *p* value <0.05 are shown.

A large majority of circadian genes have similar rhythms of gene expression in both layers. We identified and compared genes with circadian population rhythms in both human skin layers using differential rhythmicity analysis [30]. *Population* rhythms are circadian patterns of gene expression averaged across the entire cohort. We identified 1053 circadian genes in dermis and 1352 in epidermis (FDR <0.05 and amplitude >1.5 fold peak-to-trough in at least one layer, see Materials and Methods for details; lists of rhythmic genes in each layer are available as an additional file in Supplementary Table S2). 966 of these circadian genes were common to both skin layers (Figure 1C, inset, Supplementary Figure S1A). The number of circadian genes remained stable across a range of choices of FDR cutoff (Figure 1C). 386 genes were rhythmic only in the epidermis and 87 only in the dermis, as well as a further 49 genes that were rhythmic in both but with significantly different amplitude and/or phase. Thus, the expression of 917 circadian genes was indistinguishable between the two layers.

Circadian gene amplitudes are larger in the epidermis, but core clock genes are remarkably similar in the two layers. We observed a bimodal distribution of phases of all circadian dermal and epidermal genes, with peaks clustering at 1 h and 13 h after mid-sleep time on free days (MSFsc) (Figure 1D and Supplementary Figure S1A). Despite the similarity of the distributions, amplitudes of individual genes varied between layers. Epidermal circadian genes oscillated with a higher amplitude than those in the dermis among the differentially-rhythmic genes and with a trend in this direction among the circadian genes with similar rhythms (Figure 1E). There was no systematic trend in the phase difference between circadian genes common to the two layers (Figure 1F). The core clock genes were remarkably consistent (statistically indistinguishable) in amplitude and phase between the two layers (Supplementary Figure S1B) with the exception of *ARNTL* and *PER3*, which had higher amplitudes in the dermis and epidermis, respectively. Note, *NR1D1* and *NR1D2* were rhythmic in both layers but with amplitudes just outside the amplitude cutoff in the epidermis.

The similar and dissimilar rhythms in the two layers are primarily involved in cellular signaling. Circadian genes in the dermis were enriched for immune response-related pathways; in particular, multiple analyses (Reactome, KEGG, MSigDB) revealed enrichment of IL-4 and IL-13 signaling pathways (Figure 1G). However, circadian genes in the epidermis did not show any statistically significant enrichment. The differentially-rhythmic genes (circadian genes with dissimilar rhythms in the two layers) were primarily involved in the three pathways: platelet activation & signaling, signal transduction and signaling by receptor tyrosine kinases (Figure 1H).

Population circadian rhythms are not affected by choice of time reference in this dataset. We repeated the above analysis without correcting the sample collection times for subject chronotypes using MSFsc. Interestingly, the circadian genes determined with respect to external time (Supplementary Figure S2) did not deviate appreciably in number, amplitude or phase from the circadian genes (compare Figure 1C,D,E,F and Supplementary Figure S2) with respect to internal time, i.e, controlling for MSF_sc_ did not affect rhythmicity analysis significantly in our dataset.

In summary, we observed significant similarity in the circadian gene expression across these two adjacent skin layers complemented by some layer-specificity of circadian rhythms.

### Amplitudes and phases of circadian genes are subject-specific, while magnitudes are layer-specific

The population (mean) circadian gene expression describes rhythmic gene expression at the level of the cohort. We quantified next how much circadian parameters of the rhythmic genes varied among individuals in the cohort. We present this variability within the cohort in relation to the variability across layers. We fit linear-mixed-effect models [31] followed by error propagation to obtain both the average circadian gene expression (fixed effects) and the variation across subjects and layer (random effects). This analysis was performed only on the 1439 genes that showed population rhythms in at least one layer. We computed the variability in circadian parameters (magnitude (MESOR), amplitude and phase) of individual rhythmic genes across both layers and subjects (available as an additional file in Supplementary Table S3).

Magnitudes vary more across layers, and amplitudes and phases more across subjects. The error propagation analysis produced estimates of the absolute variability (as standard deviations) of the circadian parameters across subjects and layers. When the variability is viewed in absolute terms (Figure 2A), magnitude of circadian genes varied more across layers than across subjects, while amplitudes and phases were more variable across subjects than across layers. In order to better quantify the relative contribution of layer and subject to the circadian rhythm variability, we defined the fraction of variance explained by subject as 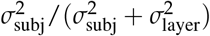. More genes were variable in amplitude and phase across subjects (have fraction of variance explained closer to one in Figure 2B) consistent with Figure 2A. However, specific genes were more or less subject-(layer-) variable in all three circadian parameters. *PER1,2* and *NR1D2* showed highly variable magnitudes and amplitudes across subjects, while positive regulators *NPAS2, ARNTL* were very consistent across subjects. The phases of core clock genes were also much more variable across subjects than across layers.

**Figure 2:**
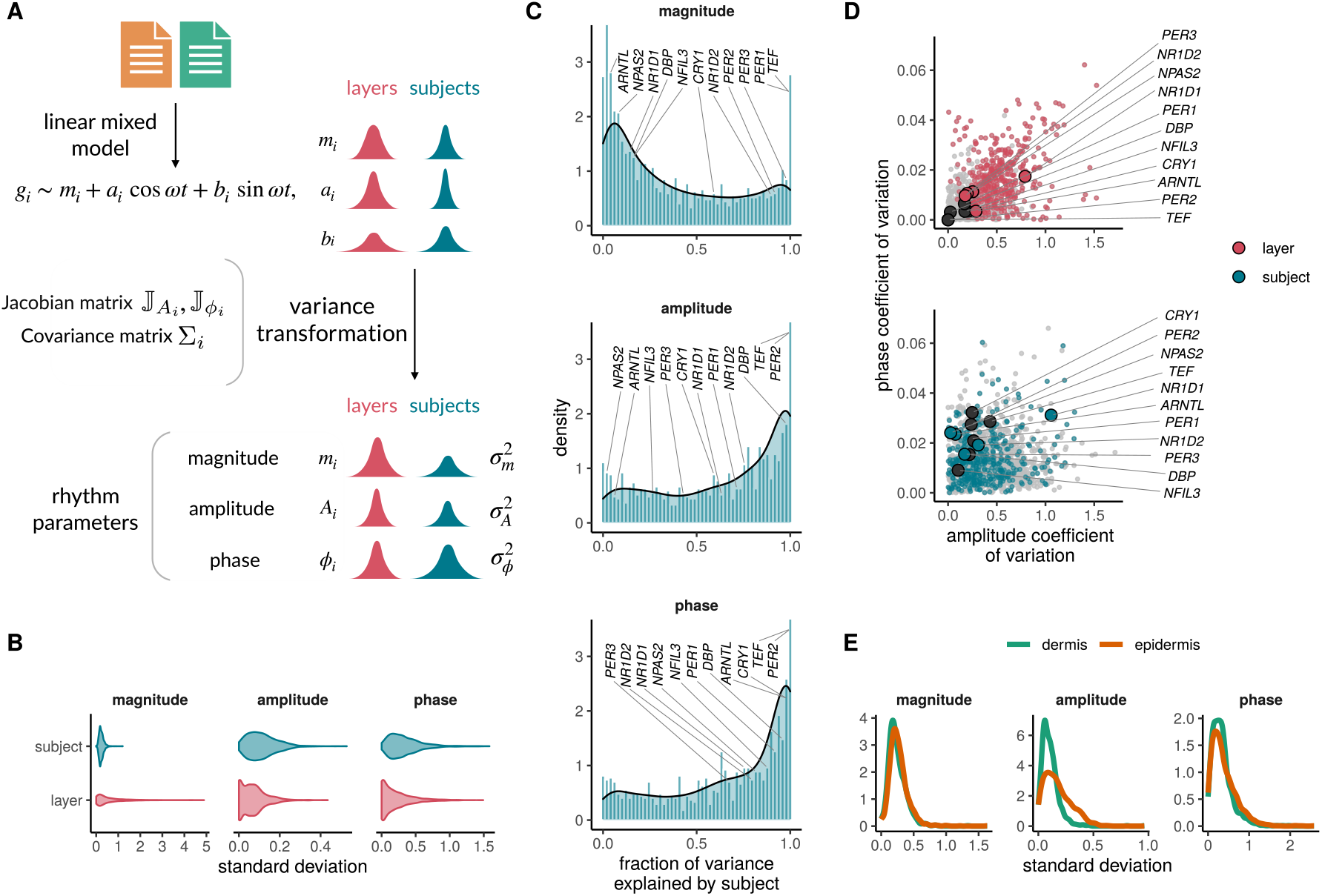
Amplitudes and phases of circadian genes differ across subjects, while magnitudes differ across skin layers. **A.** Methodology to compute variability in circadian parameters across layers and subjects. **B.** Quantification of the variability in magnitude (MESOR), amplitude and phase across subjects (blue) and layers (red) in genes (1439) rhythmic in at least one layer. **C.** Relative contribution of layer and subject to variation in magnitudes, amplitudes and phases. The fraction of variance explained by subject was defined as 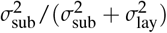 Circadian genes falling in bins closer to 0 represent cycling genes with low fraction of variance attributed to subject differences (and high fraction of variance explained by layer differences). **D.** Correlations of amplitude and phase relative variability (using the coefficient of variation) across layers (left panel) or subjects (right panel). Clock genes are shown with larger dots. Differentially-rhythmic genes (from the previous analysis) across layers are shown in color (red, blue), while genes with non-significant rhythm differences in dermis and epidermis are shown in grey. **E.** Quantification of the variability in circadian parameters across subjects in each skin layer separately. Linear mixed models were fit in all 1439 circadian genes in at least one layer to plot panels B-D; in all 1053 rhythmic genes in dermis and 1352 in epidermis to plot panel E.

Amplitudes of circadian genes are more variable than phases. The relative variation of amplitudes exceeded relative variation of phases both across layers and across subjects (Figure 2C). Genes with different population rhythms between layers (Figure 1) showed high amplitude and phase variability across layers as expected. Core clock genes had remarkably low variability in amplitude and phases across layers consistent with indistinguishable population rhythms we observed across layers. *NR1D1* was the only exception that showed larger variation in amplitude across layers; it was rhythmic in both layers but with an amplitude just below our cutoff in epidermis. However, the core clock genes had higher variability in phase across subjects, but with comparable amplitude variability across subjects and layers.

Amplitudes of circadian genes in the epidermis differ more than in dermis. We observed previously that epidermal population rhythms had greater amplitudes than dermal rhythms. In addition, when the variability was quantified in the two layers separately, amplitude variability in the epidermis across subjects exceeded that in the dermis (Figure 2D). Thus, amplitudes in the dermis were smaller but more consistent than amplitudes in the epidermis, which were larger and more variable. Nevertheless, magnitude and phase variabilities were similar in the two layers.

### Predictive biomarkers of internal time in human dermis and epidermis

Finally, we assessed the viability of skin samples to be used for circadian phenotyping. Expression of biomarker genes in the skin can serve as predictors of internal phase of entrainment or chronotype, if a fixed phase relationship between the skin clock and the central SCN clock can be assumed. To predict *internal* time from a single sample, we identified biomarkers among genes expressed in either layer individually (as suggested by the layer-specificity) using `ZeitZeiger` [32].

A small set of population rhythmic genes accurately predict internal time. For an optimal parameter choice (Supplementary Figure S3A), 8-12 rhythmic genes were sufficient to predict internal time with a median absolute error (MAE) of ~ 1.2h (Figure 3A). The ability to infer internal time from a single sample can be seen in the counter-clockwise arrangement of samples projected on the two sparse principal components (Figure 3B). The biomarkers found in the two layers all exhibited robust population circadian rhythms (Supplementary Figure S3B,C) with the exception of one gene *FOCAD*, which was also rhythmic but with an amplitude below our cutoff. The genes chosen as biomarkers all showed particularly low variability in amplitude, magnitude and phase across subjects according to our analysis in the previous section (Supplementary Figure S3D).

**Figure 3:**
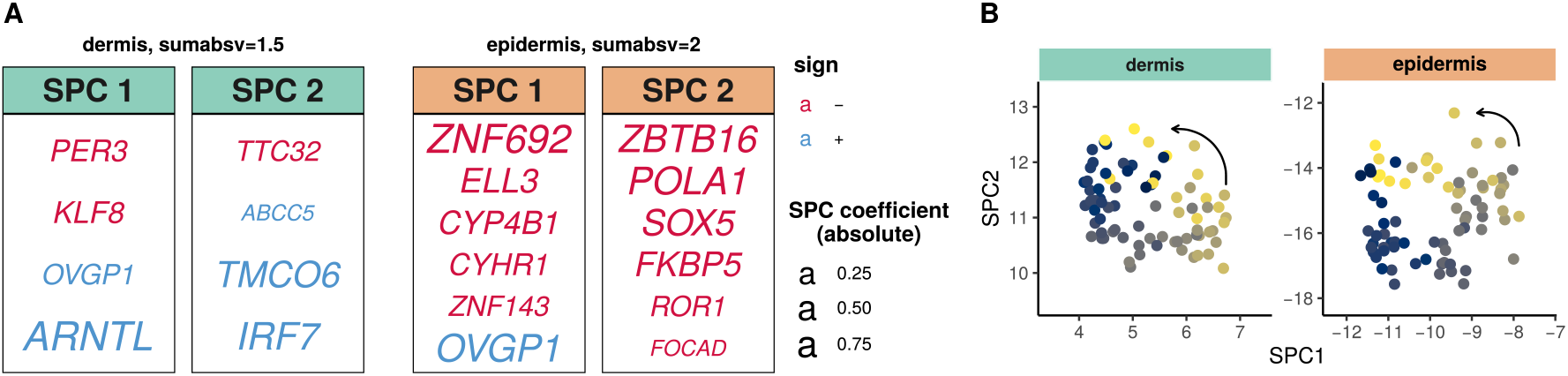
Identification of internal time-telling genes in human dermis and epidermis. **A.** Predictive biomarkers from human dermis (left panels) and epidermis (right panels) for optimal parameter choice (Supplementary Figure S3A). Genes assigned to SPC1 or SPC2 as well as their coefficients are shown. **B.** Expression profiles of the predictor genes in our cohort in dermis (left) and epidermis (bottom) represented in SPC space. Colors indicate the internal time of the subjects. ZeitZeiger was used to identify biomarkers of internal phase and was run with all ~ 11000 expressed genes, separately for dermis and epidermis.

The biomarkers sets contain layer-specific genes, but are depleted of core clock genes. Curiously, the biomarkers for internal time included only two canonical clock genes (*PER3*, *ARNTL*) and that too only in the dermal set. Moreover, the smaller biomarker set for the dermis shared only one circadian gene (*OVGP1*) with the larger set for the epidermis. The biomarkers in the dermis consisted of genes that were circadian (at the population level) in both layers. However, the epidermal biomarker set included several genes that were rhythmic only in the epidermis (*POLA1, ROR1, SOX5, ZNF143*). These sets also overlapped poorly with previously published biomarkers for epidermis (*ZBTB16, FKBP5*, [19]) and dermis (*PER3, ARNTL*, [20]).

Taken as a whole, our analysis indicates that either dermis or epidermis can be used for phenotyping circadian phase using a small set of mostly skin-specific circadian genes.

## III. Discussion

This study aimed to characterize circadian gene expression in human skin. Human studies have to cope with the heterogeneity of individuals and their clocks [25], in contrast to circadian gene expression studies in mice [33]. We addressed this challenge in several ways: First, our study only included young, healthy subjects with intermediate chronotypes and stable natural sleep-wake rhythms; we nevertheless included male and female subjects to meaningfully describe circadian rhythms in the population. Second, we corrected sampling times for chronotype differences between subjects (using mid-sleep time on free days) to present results with respect to internal time; this was only possible due to the chronobiological profiles included in our study design. Third, we structured our characterization to describe (a) the circadian gene expression on average in a random healthy member of the population and (b) extent of the deviation of the circadian gene expression of that random member from the average circadian expression.

We found ~ 1200 genes with population circadian rhythms in either layer, thus, significantly expanding on the list of known clock output genes in the skin [19, 20]. This represents ~ 10% of the expressed genes like in most circadian mammalian tissues [6, 7] suggesting that our results capture most of the rhythmic genes. This improvement resulted from the higher frequency of sampling (every 4h) in our study in relation to past studies [18–20]. However, two-thirds of these genes had indistinguishable rhythms between the layers. On the one hand, this is unsurprising given the physical proximity between the layers. On the other, it is unexpected given the well-documented heterogeneity of the skin [34] and tissue-specificity of circadian programs in physiology [7]. Core clock gene expression too was consistent between the layers with rare exceptions. Despite this general similarity, a third of the circadian genes did display expression in only one or the other layer. Epidermal circadian genes were more in number and tended to have higher amplitudes than the dermal circadian genes (as suggested previously [20]). This might be due to the greater cellular heterogeneity of the dermis or the direct exposure of the epidermis to the external environment.

To quantify the deviations of subjects from the average circadian expression, we developed a novel method based on linear-mixed models and error propagation. Unexpectedly, the magnitude of circadian genes overall remained consistent across subjects in each layer, even though it varied across layers. This result supports the claim that circadian rhythms in humans can be constructed from population sampling, i.e., one sample per individual and thus side-step cumbersome and potentially unethical longitudinal sampling [7, 19, 35]. However, amplitudes and phases of circadian genes varied more across subjects than layers (in absolute terms and as a fraction of variance). But, the expression of specific circadian genes were relatively more or less subject-variable. For instance, negative core clock members (*PER1, PER2*) had highly subject-variable magnitudes and amplitudes, while positive core clock members (*ARNTL*, *NPAS2*) were the opposite. One consistent feature of the core clock genes was the high subject-variability of their phases in both layers. This might reflect the fact that the MSF_sc_ does not fully account for the chronotype differences between the subjects. However, amplitudes generally varied more than phases measured relative to population means across subjects in each layer. Amplitudes of circadian rhythms are expected to decrease with age [36], but were not previously known to vary more than phases in similarly-aged young subjects. Finally, dermal rhythm amplitudes were smaller but less variable across subjects, while epidermal rhythms possessed larger amplitudes and were more subject-variable. This represents an interesting dichotomy: the dermis might be a better source of circadian biomarkers, but the epidermis might be more indicative of amplitude differences between individuals with a larger dynamic range.

In recent years, a number of novel approaches have been introduced to assess circadian parameters and, in particular, circadian phase in humans (see [37] for a nice review) based on machine-learning on high-dimensional -omics data. We therefore explored the suitability of these two skin layers to provide biomarkers to predict internal clock phase from single samples. Similar to our previous work on blood-based circadian phase determination [38], the expression of a small set of 8-12 circadian genes at a single time point was sufficient in either skin layer to predict internal clock phase with a median accuracy of ~ 1 h. This accuracy is probably optimistic as it is based on internal cross-validation and a separate validation is necessary to estimate its true accuracy. Even with some loss of accuracy these biomarkers might be expected to perform as well as biomarker sets previously proposed for the epidermis and dermis [19, 20]. Genes in biomarker sets must possess consistent magnitudes and amplitudes across subjects in addition to robust rhythmicity in order for the inference from a single sample to be feasible. Our biomarker set for each layer showed low magnitude and amplitude variability across subjects as is desirable. Unlike other identified circadian biomarkers for phase [19, 20, 38, 39], the sets discovered in this study were almost devoid of core clock genes. Moreover, most biomarker genes were either rhythmic only in the layer in question or the genes differed significantly in amplitude and/phase from the other layer. This raises the unexpected possibility that biomarker sets involving tissue/layer-specific circadian genes might be better suited for internal phase prediction than core clock genes.

Our results leave open questions that need to be addressed in future studies. Our cohort was small, healthy, young and Caucasian and it is unclear how much our results can be extrapolated to a diverse population. Our inability to find differences between an analysis based on internal and external time was surprising, but is probably due to both the lack of extreme chronotypes in our cohort and insufficiency of 4 h sampling resolution to accurately reflect the ~6h range of 95% of human chronotypes. The estimates of the variability in gene expression across subjects are affected by the small cohort size. In fact, the inter-subject variability subsumes inter-sex variability and cannot be reliably separated in this small cohort. Variance estimates across just two layers are well defined, but are likely less accurate compared to variance estimates across subjects. The error propagation analysis to quantify variation of circadian parameters is based on linearization and hence, assumes estimated mixed effects are “small”. We identified circadian biomarkers for healthy young individuals maintaining natural yet regular sleep schedules. Whether these are also good markers in elderly and sick individuals and those with disrupted sleep schedules, such as shift workers, remains unanswered. It is unknown whether the human skin clock has a fixed phase relationship too or is independent of the central clock [40]. If the former, then the identified biomarkers can be used to predict central clock phase, else they would only be able to predict peripheral skin clock phase.

## IV. Conclusions

We presented the most complete description to date of the transcriptional output of the circadian clock in two human skin layers. We reported both how average gene expression rhythmically varies in the population and the inherent variability in these average rhythms due to population heterogeneity. Our consideration of internal time in the analysis makes our results applicable to humans regardless of chronotype. Not only does our work provide a comprehensive resource of circadian gene expression and circadian biomarkers for phase in human skin, but also provides a methodology to describe human circadian rhythms in a population.

## V. Materials and Methods

### Experimental design and collection of human skin samples

The study to obtain human skin biopsies was approved by the local Ethical Review Board at Charité Universitätsmedizin Berlin (EA4/019/11). Tissue samples were collected according to the recommendations of the Declaration of Helsinki and to applicable laws for a non-drug study. All donors provided written and informed consent. The study was performed at the Clinic for Sleep & Chronomedicine, St. Hedwig-Krankenhaus Berlin. Eleven healthy volunteers (six males, five females, aged 20-30 years in 2011) participated in the study. Individual chronotypes were assessed using the Munich Chronotype Questionnaire (MCTQ) by calculating the mid sleep time on free days adjusted for the sleep-debt accumulated during the workweek MSF_sc_ [29]. All information about the subjects is provided in Supplementary Table S1.

Three millimeter biopsies were obtained from the upper back for seven time points over a period of 24 h (8:00, 12:00, 16:00, 20:00, 00:00, 04:00 and 8:00 the following day). Skin biopsies were subsequently incubated in PBS at 55 °C for 3 min to separate epidermis and dermis. Tissue samples were then frozen in liquid nitrogen and stored at −80 °C. RNA extraction and quality control from skin biopsies was performed by Miltenyi Biotec using the TRIzol method. Linear amplification and labeling of RNA and hybridization of Agilent Whole Genome Oligo Microarrays 4×44k (Agilent Technologies) using 1.2–1.65 μg of Cy3-labeled cRNA was performed by Miltenyi Biotec, essentially as reported in [41].

### Gene expression analysis

The microarray gene expression analysis was conducted in R. The RMA (Robust Multichip Average) algorithm was used to pre-process and extract expression profiles from the raw CEL files. Genes were annotated with ENSEMBL and ENTREZ IDs using Agilent “Human Genome, Whole” annotation data (hgug4112a.db, v3.2). The raw gene expression data has been deposited in the Gene Expression Omnibus (GEO) under the accession number GSExxxx.

Raw data of the hybridized microarrays were normalized and processed using the Bioconductor R-Project package Linear Models for Microarray Data (`limma`). Background correction, normalization between the different arrays, removal of non-annotated probes, lowly-expressed genes as well as control probes was performed as suggested by the limma user-guide. The filtered and annotated dataset contains 11578 expressed genes.

Principal Component Analysis (PCA) was performed in order to remove outliers. The expressed genes nicely organized in two clusters in PCA space, separated by skin layer (data not shown). Nevertheless, the epidermis sample from subject 109 taken at 8:00 the following day did not cluster with the rest of epidermal samples and for this reason was removed from further analyses.

### Rhythmicity analysis and functional annotation of circadian gene lists

To detect genes exhibiting rhythmic behavior with a 24 h period in their expression we used cosinor analysis and differential rhythmicity analysis as decribed in [30]. In short, we first tested the null hypothesis that the sine and cosine terms from cosinor analysis are equal to 0. Acrophases and amplitudes were estimated from the analysis and used to identify significantly oscillating genes. If the null hypothesis could be rejected under a false discovery rate threshold < 0.05 and a minimum amplitude requirement (i.e., that either the amplitude of the oscillating gene in dermis *or* in epidermis is above a peak-to-trough fold change amplitude >1.5), we classified that gene as rhythmic in at least one of the layers. Next, we tested the differential rhythmicity null hypothesis, namely that, among the genes that were rhythmic in at least one layer, sine and cosine terms are equal across skin layers. If this hypothesis could be rejected under a FDR< 0.05, the gene was considered to have significantly *different* rhythms in dermis compared to epidermis. If, on the contrary, the null hypothesis could not be rejected, we defined the gene to have *indistinguishable rhythms* across layers. Importantly, genes with amplitudes below the threshold in one of the layers but above the threshold in the other layer (that passed the minimum amplitude requirement) and with statistically indistinguishable rhythms were considered rhythmic and were included in our analyses despite the amplitude value being below the threshold in the first step of the analysis. Lists of rhythmic genes together with their amplitude and phase values are available as an additional file in Supplementary Table S2.

In analyses where *internal* time was used, sampling (wall) time was corrected to internal time in each subject by subtracting the mid-sleep time on free days after correcting for sleep debt during week days (MSF_sc_ [29]) to wall time.

### Assessment of variability in circadian parameters across subjects and skin layers

In order to analyze how magnitudes, amplitudes and phases of individual circadian genes vary across subjects and layers, we analyzed each gene separately using the linear mixed models [31, 42, 43]. The expression of gene *i, g_i_*, is modeled as

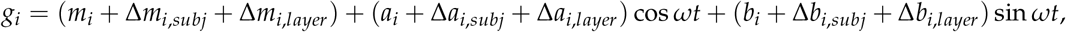

where *m_i_, a_i_* and *b_i_* represent the coefficients of the fixed effects for gene *i*; and Δ*m_i_*, Δ*a_i_* and Δ*b_i_* represent the random effects attributed to differences across layers or subjects, which are drawn from a normal distribution, whose variance is estimated.

While *m_i_* is a direct readout of the gene’s magnitude (and its respective uncertainty attributed to layers/subjects), amplitude *A_i_* and phase *ϕ_i_* of a gene *i* were calculated from the coefficients *a_i_* and *b_i_* for each gene as 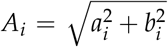 and 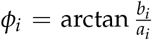. To determine the variability in amplitude and phase across subjects and layers, we used error propagation. The variance of amplitude and phase across subjects and layers (*x* = *layer*,*subj*) was computed from the Jacobian matrices 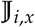 and the covariance matrices *Σ_i,x_* of the rhythm parameters (obtained from the linear mixed-model fits) as

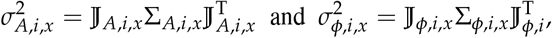

### Identification of predictive biomarkers of molecular skin phase

We used ZeitZeiger [32] to identify skin biomarkers of circadian phase. We tested two sets of predictors using the whole set of expressed genes in epidermis or dermis separately. The predicted variable was, in both cases, internal time. To evaluate the performance of the predictors, we followed a leave-one-subject-out cross-validation approach in the lines of [32, 38]. To do this, predictors are trained with data from all subjects except one and internal time from the subject who is left-out is predicted. The process is iterated along all subjects and for different values of the two main parameters of ZeitZeiger, `sumabsv` and `nSPC`. The first parameter `sumabsv` controls how many genes form each sparse principal component (SPC) and the second parameter, `nSPC`, controls how many SPCs are used for prediction. Large values of either parameter result in more genes being needed for prediction. For each set of values of `sumabsv` and `nSPC` from the leave-one-subject-out cross-validation, we calculated the median absolute difference between the predicted and the observed internal time stamp across all subjects. We refer to this parameter as median absolute error (MAE), and it serves as a measure of accuracy of the prediction: the lower the error, the better the prediction.

## Abbreviations

SCN: suprachiasmatic nucleus
MSFsc: mid sleep time on free days after correcting for sleep debt during week days
FDR: false discovery rate
MAE: median absolute error
GEO: Gene Expression Omnibus
SPC: sparse principal component.

## Conflict of interest

AK received consulting fees from Beiersdorf AG. The rest of the authors have on conflict of interest to declare.

## Data accessibility

The gene expression microarray data of dermis and epidermis generated in this study are available at the Gene Expression Omnibus (GEO) database and can be accessed under the accession number GSExxxx. The source code is available through GitHub: xxxx.

## Funding

This study was supported by Deutsche Forschungsgemeinschaft (DFG, German Research Foundation) grant AN 1553/2-1 to BA and by the Project-ID 278001972 – TRR 186 to HH, AK and MdO.

## Author contributions

Study design and conceptualization: BA, MdO, AK, DK, HH, HR and TB. Experimental Methodology: SK, FS, AG, JdZ and CN. Bioinformatic methodology: MdO, BA and KJ. Investigation: BA, MdO, KJ, AK and HH. Writing (original draft and editing): MdO and BA. Writing (review): MdO, BA, MF, AK, HH, HR, TB, MF and JdZ. Funding acquisition: BA, AK, TB and HH.

## Notes

### Summary of Updates

Funding information revised; new supplementary table including list of genes with population circadian gene expression in each layer (Supp. Table S2); included captions for external supplementary tables.

