## Supplementary Information for "Inter-layer and inter-subject variability of circadian gene expression in human skin"

### Supplementary Material

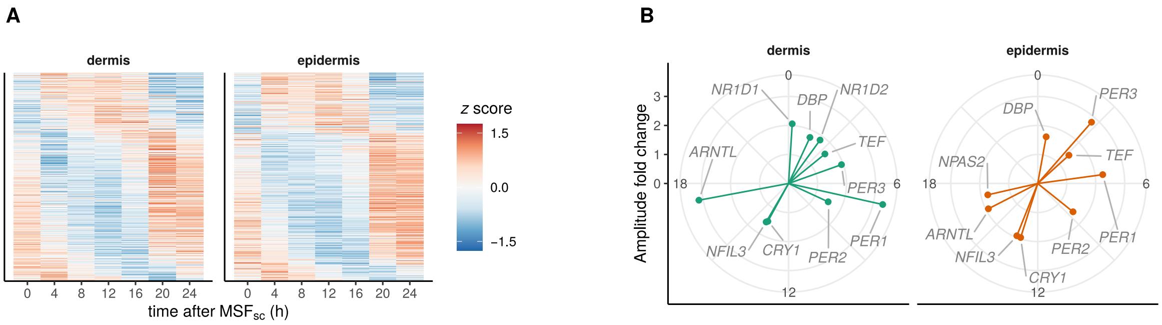


**Figure S1: Population circadian gene expression in healthy human skin. A.** *z* score-normalized, acrophase-ordered expression heatmap of the circadian genes from human dermis (left) and epidermis (right). **B.** Expression profiles of circadian core clock genes in human dermis (left) and epidermis (right). Arrow direction represents phase (expressed as peak time, in hours, after MSF$_{sc}$) and arrow length depicts peak-to-trough fold change amplitude.

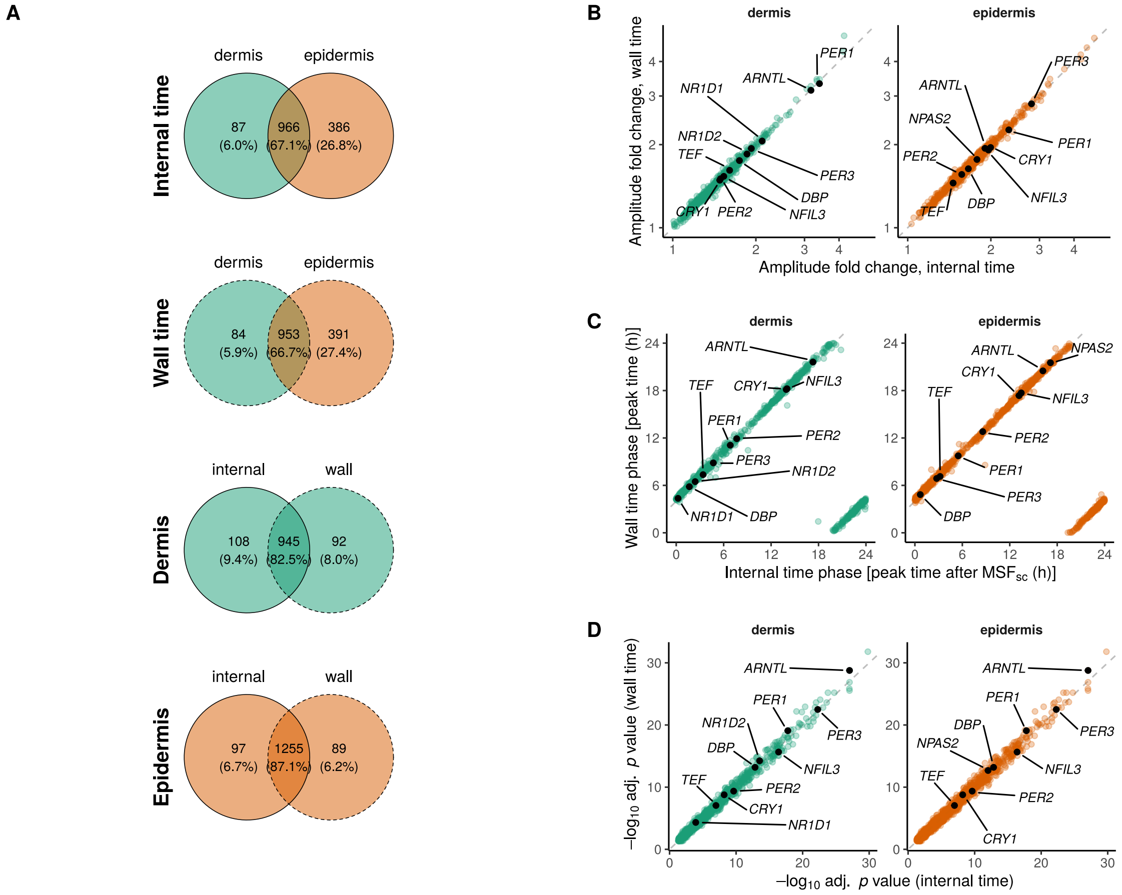


**Figure S2: Population circadian rhythms in human skin are similar when time is not adjusted for chronotype differences. A.** Venn diagram visualization of the number of genes identified as circadian in dermis (green) vs. epidermis (orange) and in the analysis using internal time (i.e., after correcting for chronotype differences, solid line) or wall time (dashed line). **B.** Amplitude correlation of genes identified as rhythmic with the internal time analysis compared to external time analysis. **C.** Phase correlation of circadian genes identified with the internal time analysis compared to external time analysis. Phase with respect to internal time was defined as peak time after $MSF_{sc}$, while phase with respect to wall time is the peaking time of the respective gene without correction. **D.** Benjamini Hochberg-adjusted $p$ value correlation of rhythmic genes identified with internal vs. external time analysis.

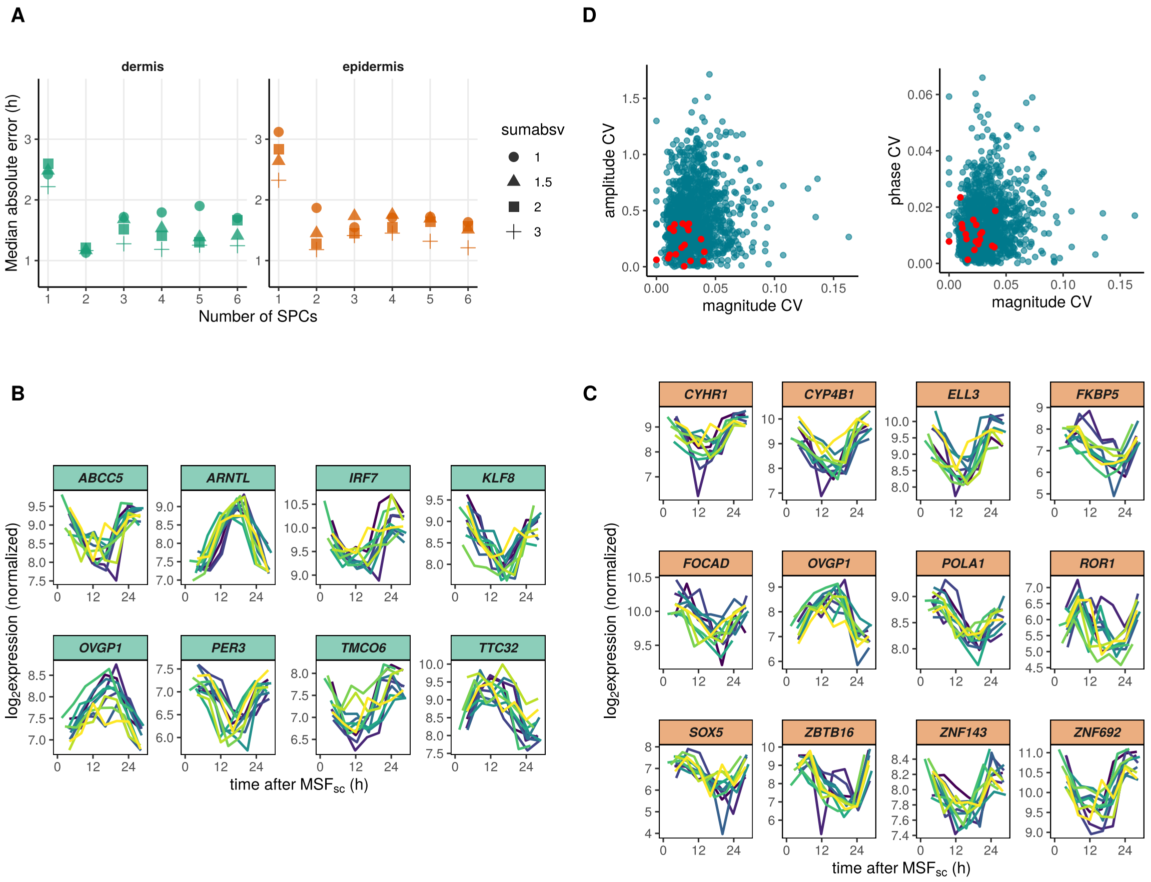


**Figure S3: Predictive biomarkers of internal time in human dermis and epidermis. A.** Median absolute error of the internal-time prediction on cross-validation (see Materials and Methods for details) as a function of the two main parameters of `ZeitZeiger`, `sumabsv` and `nSPC`. **B.** Quantification of magnitude, amplitude and phase variability of circadian genes across subjects. Predictive biomarkers are shown in red. **C.** Expression profiles of the time-telling genes in dermis (green) and **D.** epidermis (orange) for optimal parameter choice of `sumabsv` and `nSPC`. Colored lines represent the time series in different subjects. `ZeitZeiger` was run with all $\sim$ 11000 expressed genes, separately for dermis and epidermis.

**Table S1: Information and sleeping schedules of the healthy subjects who participated in the study.** Chronotypes were estimated from sleep schedules as the mid-sleep time on free days after correcting for sleep debt ($MSF_{sc}$) [1, 2].

| Subject | Sex | Birth year | Bed time work days | Sleep time work days | Min fall asleep work days | Wake up time work days | Min wake up work days | Alarm work days? |
|---|---|---|---|---|---|---|---|---|
| P108 | male | 1984 | 22:45 | 23:00 | 20 | 6:30 | 15 | Y |
| P100 | male | 1991 | 23:00 | 23:00 | 7.5 | 7:00 | 5 | Y |
| P113 | male | 1985 | 23:30 | 0:00 | 30 | 8:30 | 10 | Y |
| P106 | male | 1982 | 23:00 | 23:15 | 15 | 6:52 | 7.5 | Y |
| P102 | male | 1989 | 23:00 | 23:00 | 10 | 7:30 | 0 | Y |
| P109 | male | 1984 | 23:00 | 23:00 | 5 | 8:00 | 0 | Y |
| P103 | female | 1988 | 0:00 | 0:25 | 25 | 7:30 | 9 | Y |
| P107 | female | 1987 | 23:00 | 23:00 | 15 | 6:00 | 5 | Y |
| P111 | female | 1983 | 0:00 | 0:00 | 5 | 8:00 | 15 | Y |
| P114 | female | 1986 | 23:30 | 23:30 | 5 | 8:30 | 5 | Y |
| P115 | female | 1981 | 22:00 | 22:15 | 5 | 5:20 | 5 | Y |

| Subject | Bed time free days | Sleep time free days | Min fall asleep free days | Wake up time free days | Min Wake up time free days | Alarm free days? | Corrected mid sleep time |
|---|---|---|---|---|---|---|---|
| P108 | 23:00 | 23:15 | 15 | 6:30 | 30 | N | 02:58 |
| P100 | 0:00 | 0:00 | 5 | 8:00 | 15 | Y | 04:00 |
| P113 | 1:00 | 1:15 | 20 | 9:30 | 15 | Y | 05:28 |
| P106 | 23:30 | 23:45 | 15 | 9:15 | 60 | Y | 03:50 |
| P102 | 0:00 | 0:00 | 10 | 8:00 | 10 | N | 04:11 |
| P109 | 0:00 | 0:00 | 5 | 8:30 | 30 | N | 04:26 |
| P103 | 0:00 | 0:00 | 25 | 7:30 | 5 | Y | 03:36 |
| P107 | 0:00 | 0:00 | 15 | 7:30 | 5 | N | 03:34 |
| P111 | 2:30 | 2:30 | 5 | 11:00 | 30 | N | 06:34 |
| P114 | 23:30 | 23:30 | 5 | 8:30 | 5 | N | 04:00 |
| P115 | 0:00 | 0:15 | 5 | 8:30 | 30 | N | 03:58 |

**Table S2: List of genes with population circadian gene expression in each layer.** External .xlsx file.

**Table S3: Estimated variability of circadian gene expression parameters across subjects and layers.** External .xlsx file.
